# Endogenous antimicrobial peptides in human colostrum: a peptidomics study by liquid-chromatography high-resolution mass-spectrometry (LC-HRMS) and bioinformatics

**DOI:** 10.1101/2025.02.03.636292

**Authors:** Isabele B. Campanhon, Nicole Cavalcante da Silva, Rosane de Oliveira Nunes, Flávia Fioruci Bezerra, Alexandre Guedes Torres, Márcia R. Soares

## Abstract

Human colostrum is recognized as a source of bioactive compounds that impact on neonatal development, such as antimicrobial peptides. Antimicrobial peptides (AMPs) present some common characteristics, as they are relatively small (<10 kDa), most of them are positively charged and possess α-helical regions. These molecules can protect the newborn against infectious agents such as bacteria, fungi, and viruses. Using an MS-based proteomic approach combined with the application of bioinformatic tools, 29 peptide sequences from human colostrum were annotated in human colostrum. These peptides are potentially antifungal and were enriched in glutamic acid (E), phenylalanine (F), lysine (K) and tryptophan (W) residues. Among the precursor proteins, two have a known 3D structure and their fragments are located in the protein core. In addition, *in silico* analyses revealed that AMPs containing regions of α-helix and random structures, and high aliphatic index (mean of 85.29) tend to be thermostable. In this work with human colostrum, previously unreported peptides with potential antimicrobial activity were annotated and structurally characterized via bioinformatic tools.

## 1. Introduction

The main causes of neonatal death include infections, prematurity and birth asphyxia, and the protective role of breast milk against infections has been widely reported [1–3]. Protective factors in breast milk include immunoglobulins, and bioactive compounds, such as antihypertensive, antioxidant, antithrombotic and antimicrobial peptides [9]. These factors can also stimulate the maturation of the neonatal immune system [4,5]. Antimicrobial peptides (AMPs) were active against several pathogens [10] and thus they might reduce neonatal mortality by infections. Endogenous peptides can be released via the action of milk proteases such as plasmin, cathepsin D and elastase, and proteolytic activity can be modulated by activators and inhibitors secreted by mammary epithelial cells [6]. Some of these endogenous proteases may be active in the mammary alveoli before milk ejection [7], and the extent of proteolytic activity between these proteases might differ in intact milk and during infant digestion [8].

Antimicrobial peptides in breast milk might protect the newborn against emerging viral diseases, such as COVID-19, and to bacteria resistant to antimicrobials [11]. In addition, the prospection for endogenous bioactive peptides may contribute to future developments of infant formulas that would help to prevent neonatal infections [12]. As most studies on bioactive peptides in breast milk investigated those formed during *in vitro* digestion of mature milk [13,14], endogenous peptides in colostrum remain largely unresearched. Previous studies showed that the profile of endogenous peptides in milk varied between women delivering at term and those delivering prematurely, and peptides in these groups were expressed differently [15,16]. In this work, colostrum ultrafiltration followed by nano-liquid-chromatography coupled to high-resolution mass spectrometry (nLC-Orbitrap-HRMS), combined with bioinformatic tools were used to annotate and characterize the endogenous peptides of human colostrum with potential antifungal activity. Optimization of antimicrobial peptide structures might support future clinical applications, in addition to being a natural source for the discovery of new antimicrobial peptides.

## 2. Materials and methods

### 2.1. Study volunteers and breast milk collection

Mothers who delivered at term were recruited, and colostrum samples were obtained from twenty-four healthy adult volunteers. Samples were collected by manual expression into polypropylene containers at the Herculano Pinheiro Maternity Hospital in Rio de Janeiro, which is a public health service unit. All donors were between the ages of 21 and 40 years with no diagnosed chronic diseases, and were not taking any prescribed medications or vitamin supplements. Participants were recruited between 37 to 41 weeks of pregnancy, and colostrum samples were collected between 1 and 5 days postpartum by standard procedures to avoid microbial contamination. The research protocol was approved by the Ethical Committee of Maternity School of the Federal University of Rio de Janeiro (protocol number CAAE: 56617516.5.0000.5259), and the mothers signed informed consent before participating in the study. Milk samples were transported on ice to the laboratory and stored at –80 °C until analysis. Maternal and neonatal characteristics are presented in **Supplementary Table 1**.

### 2.2. Sample preparation

Colostrum samples were thawed in a 37 °C water bath and protein concentration was determined according to the Lowry [17] protein assay. Subsequently, a 5.0 mL pooled sample was prepared, containing all 24 samples balanced for their protein contents so that the pooled sample had the same amount of protein from each donor milk.

Milk fat was removed by centrifuging (1,500 ×*g*, 20 min) and collecting the defatted aqueous fraction consisting of skim milk [18]. The skim milk was collected and ultra-filtered through Microcon^®^ concentrators (YM-10K, Millipore, Billerica, MA, USA), at 14,000 ×*g* for 15 minutes at 10 °C to separate the peptide (< 10 kDa, filtrate) and protein (≥10 kDa, retained) fractions.

### 2.3. Mass spectrometry analysis and bioinformatics

The samples were analyzed in triplicate in a nano-LC Ultimate 3000 (Thermo Fisher Scientific, IL, USA) coupled to a quadrupole-Orbitrap mass analyzer platform (Q-Exactive Plus; Thermo Fisher Scientific, Bremen, GER). An aliquot containing 1 µg of peptides was loaded on a guard-column (2 cm length, 200 µm i.d.) packed with ReproSil-Pur C18-AQ (5 µm particle diameter) and separated in an analytical column Picochip with ReproSil-Pur C18 (3 µm particle diameter). Peptides were eluted using a gradient of 95% solvent A (95% H_2_O, 5% acetonitrile, 0.1% formic acid) to 40% solvent B (95% acetonitrile, 5% H_2_O, 0.1% formic acid) for 60 min; 40% to 85% of solvent B for 10 min; and 95% of solvent B for 15 min, at a 300 nL/min flow rate.

Full scan and MS/MS were acquired on positive mode using a data-dependent acquisition (DDA; dd-MS^2^) protocol. The ion source and S-lens were optimized to spray voltage at 3.4 kV, zero flow of sheath and auxiliary gas at 250 °C and 80 S-Lens RF level. MS full scan was acquired at 70,000 mass resolution scanning from 400 to 2000 m/*z* in the Orbitrap analyzer, 10^6^ AGC, and 50 ms maximum ion injection time. The 10 most intense ions containing 2 to 4 charges were selected for high-energy collision dissociation (HCD) fragmentation. Fragment acquisition in the Orbitrap using 30 a.u. normalized collision energy, and dynamic exclusion was enabled for 20 s. The MS/MS scans were obtained using 17,500 resolution, 5 × 10^4^ AGC, 50 ms maximum isolation time, 1 sec microscans, 200-2000 m/*z* range, and 2.0 m/*z* isolation window.

The DDA raw data were processed and searched by the Proteome Discovery 2.1 software server search engine (Thermo Fisher Scientific, Bremen, GER) with the SEQUEST algorithm by tolerating up to ± 0.1 Da for precursor ions and 0.01 Da for fragment ions. Protein identification was performed by searching the mass spectrometric data against the UniProt-SwissProt protein database (available in April 2018) containing reversed sequences with a false discovery rate (FDR) < 1 %.

### 2.4. Search for functional endogenous peptides

Peptide sequences obtained by mass spectrometry were compared to sequences from four functional peptide databases: Milk bioactive peptide database [19], AMPA (Antimicrobial Sequence Scanning System) [20], ClassAMP (A prediction tool for classification of AMPs) [21] and APD (The antimicrobial peptide database) [22]. The milk bioactive peptide database was constructed from peptides derived from milk proteins described in previous research articles with a biological function [19]. The AMPA tool is based on the characteristics of each amino acid and is used to classify different protein sequences with antimicrobial or non-antimicrobial regions. ClassAMP was used to predict the propensity of a peptide sequence to present antibacterial, antifungal or antiviral activity. APD was used to assess the similarity between the sequences annotated and those deposited in public libraries, and only the peptides with at least 40% similarity were considered.

### 2.5. In silico *analysis of the three-dimensional structure of proteins and peptides*

The three-dimensional structure of the precursor proteins was obtained from The Protein Data Bank (PDB), and PyMOL software [26] was used to import PDB files. For the peptides sequences, the three-dimensional structure was predicted by the software PEPFOLD3, an online resource for *de novo* peptide structure prediction [27].

### 2.6. Physicochemical properties of peptides

Molecular mass, isoelectric point, the number and composition of amino acid residues, values of the instability index, aliphatic index and GRAVY index (grand average of hydropathicity) of antimicrobial peptides were determined *in silico* by ExPASy-ProtParam (http://web.expasy.org/protparam). In addition, the amino acid frequencies were calculated by Genscript software (https://www.genscript.com/tools/codon-frequency-table).

### 2.7. Evaluation of the antimicrobial activity

Peptide sequences were classified as best AMP candidates if they were matched by at least three bioinformatic tools. Two of the most promising peptides were synthetized commercially and used to assess antimicrobial potential. *Escherichia coli* strain DH5α was purchased from New England Biolabs (NEB). The antimicrobial activity of the two peptides: AGLAPYKLRPV derived from lactoferrin and HLPLPLLQPL derived from beta-casein was evaluated on batch cultures containing 0.001, 0.01 and 0.1 mg/mL– against *E. coli* DH5alfa in a cell suspension of 10^8^ CFU/mL grown in LB media (Luria-Bertani) (BD™, Le Pont de Claix, France). Cells were then serially diluted 10-fold each step in 8.5 g/L NaCL, 2.0 g/L Tween 80, plated on solid LB and after incubation at 37 °C for 18 h the colony-forming units were counted. The procedure was performed in triplicate, and inhibition was calculated as described elsewhere [28].

### 2.8. Statistical analysis

Data were expressed as mean ± standard deviation, and were processed using Prism for Windows software, version 5.04 (GraphPad Software, San Diego, CA). Paired *t-*tests were used to test differences between peptide sequences. A *p*-value < 0.05 was considered as statistically significant.

## 3. Results

### 3.1. Maternal and neonatal characteristics

**Supplementary Table 1** shows the characteristics of the colostrum donors and their newborns during sample collection.

### 3.2. Endogenous Peptides analysis

Endogenous peptides were determined in a pooled sample of human colostrum from 24 adult mothers. Colostrum was defatted by centrifugation, the skimmed colostrum was ultrafiltered (through 10 kDa cut-off membranes), and the ultrafiltered fraction was analyzed (< 10 kDa peptides). A total of 29 peptide sequences were identified by Mass Spectrometry (MS/MS) deriving from 15 precursor proteins (**Table 1**). The major milk proteins such as caseins, lactoferrin and immunoglobulins were identified among precursors of the peptide fragments. Most (> 40 %) of the peptide sequences were derived from β-casein.

**Table 1:**
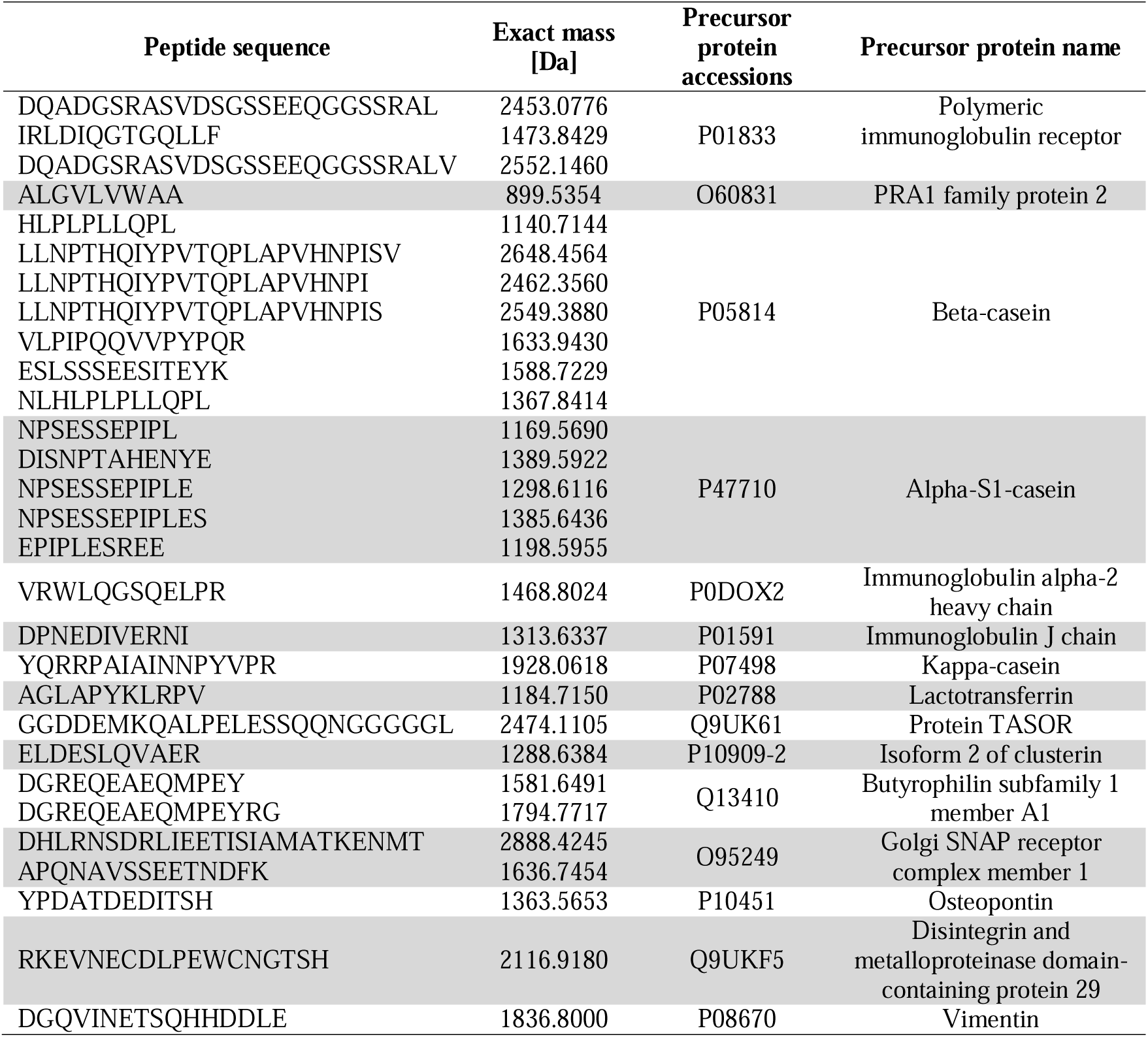
Peptide sequences in the endogenous fraction (<10 kDa) of human colostrum (pooled sample, from *n* = 24 adult donors) annotated by mass spectrometry (nLC-HRMS/MS, Orbitrap), and their precursor proteins.

### 3.3. Physicochemical properties of peptides

Physicochemical properties of individual peptides are presented in **Table 2**, showing that some of these properties varied broadly among the annotated peptides. Most of the peptides showed a negative hydropathy index value indicating that these sequences have a hydrophilic nature (**Table 2**). In the two groups of peptides showing either hydrophilic or hydrophobic nature, the isoelectric point was on average 5.5 and 5.7, respectively, indicating a mildly acidic nature. The annotated peptides contained from 9 to 24 amino acid residues, and the frequency of occurrence of each amino acid is shown in **Figure 1**.

**Fig. 1.**
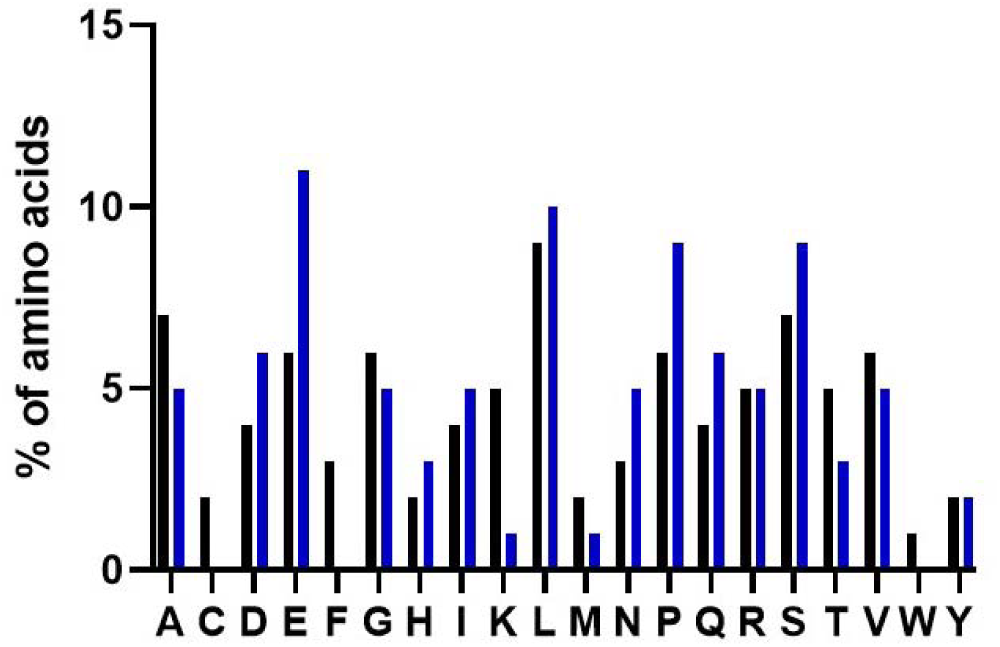
Frequency of amino acids found in the endogenous peptides’ sequences in human colostrum 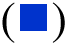 annotated by mass spectrometry (nLC-HRMS/MS Orbitrap) compared to the frequency of amino acids present in human proteins 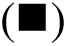 calculated using the Genscript program (https://www.genscript.com/tools/codon-frequency-table). The one-letter coding of amino acid names was used in this Figure.

**Table 2.**
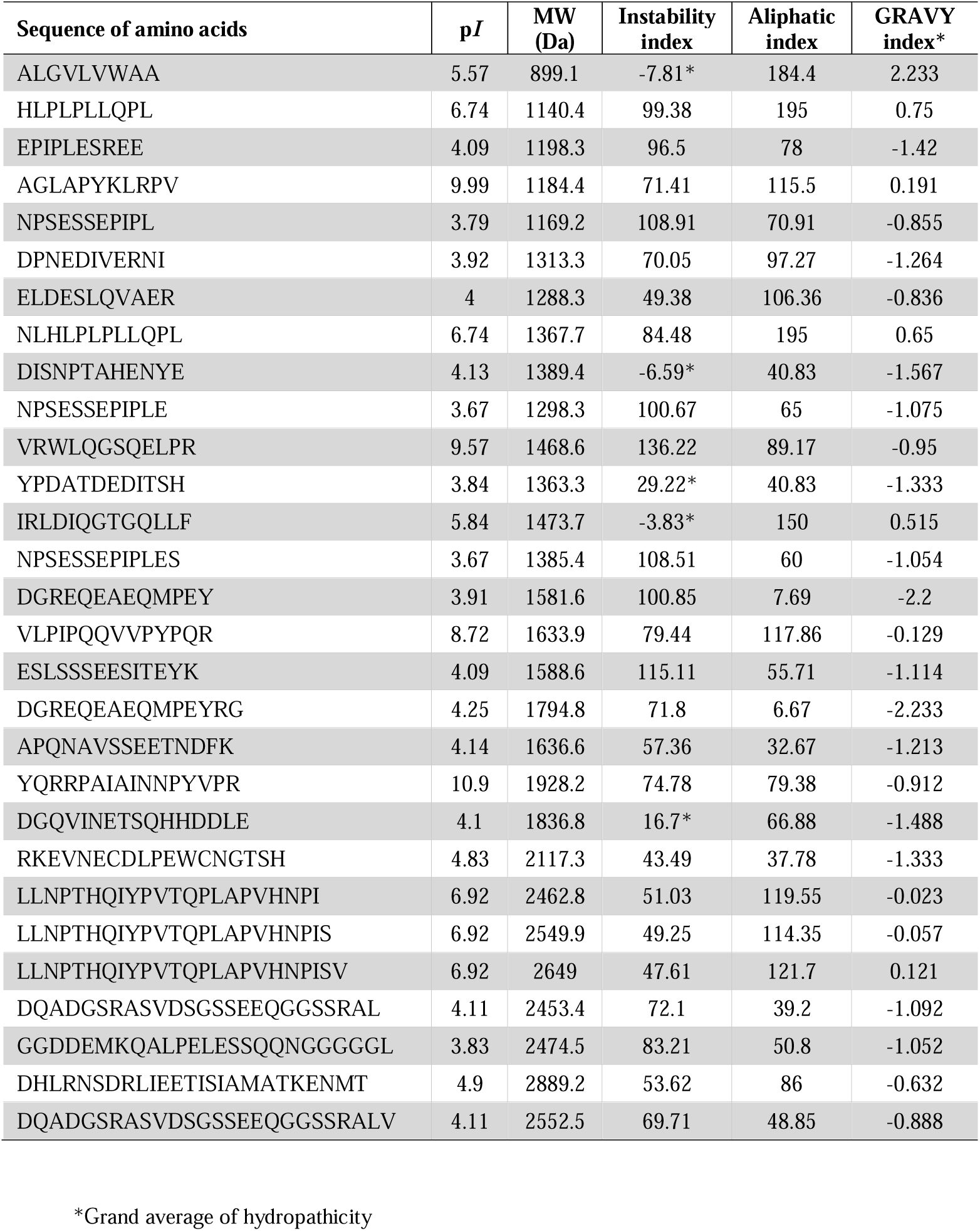
Physicochemical characteristics of endogenous peptides (fraction < 10 kDa) in human colostrum annotated by mass spectrometry (nLC-HRMS/MS Orbitrap) evaluated by ProtParam software.

Although the amino acids’ frequency did not vary significantly (paired *t*-test), the amino acids glutamic acid (E), phenylalanine (F), lysine (K) and tryptophan (W) were more prominent.

The aliphatic index of the sequences ranged from 6.67 to 195, but on average it was high (85.29), which is positively correlated with thermostability. The most stable peptides were IRLDIQGTGQLLF from polymeric immunoglobulin receptor, DISNPTAHENYE from alpha-S1-casein, ALGVLVWAA from PRA1 family protein 2, YPDATDEDITSH from osteopontin and DGQVINETSQHHDDLE from vimentin.

### 3.4. Analysis of peptides with potential antimicrobial activity

In this study, we searched four peptide databases for the annotated endogenous peptides in colostrum. The sequences identified herein have not been previously deposited in the databases of the bioinformatics service Milk Bioactive Peptide Database [19] and AMPA software [20]. In addition, the peptide sequences identified were tested in the search software ClassAMP for antimicrobial functions, and all the peptides presented potential antimicrobial activity (**Supplementary Table 2**). Among these peptides, 45 % were predicted with potential antifungal activity (**Figure 2**).

**Fig. 2.**
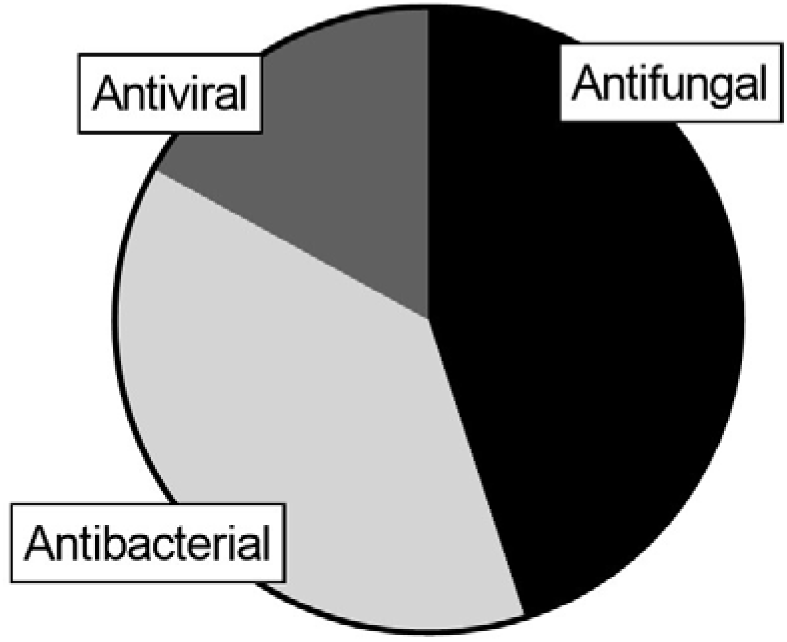
Antimicrobial activity of endogenous peptides identified by mass spectrometry (nLC-HRMS/MS Orbitrap) in human colostrum using the ClassAMP program. Antifungal activity (45%), antibacterial activity (38%), and antiviral activity (17%).

Additionally, the APD database was used to assess the similarity between the annotated peptide sequences and those deposited in the bank. APD is based on the physicochemical properties of peptides [22]. Some peptides showed a similarity near 40%, these sequences are presented in **Supplementary Table 3**.

### 3.5. In silico *3D structure prediction of endogenous peptides*

The prediction of peptides three-dimensional structure was performed using the PEPFOLD3 software [27], which allows structure prediction of peptides that have between 9 and 36 amino acids in aqueous solution. Structural models of the 29 peptides obtained by this software are shown in **Figure 3**.

**Fig. 3.**
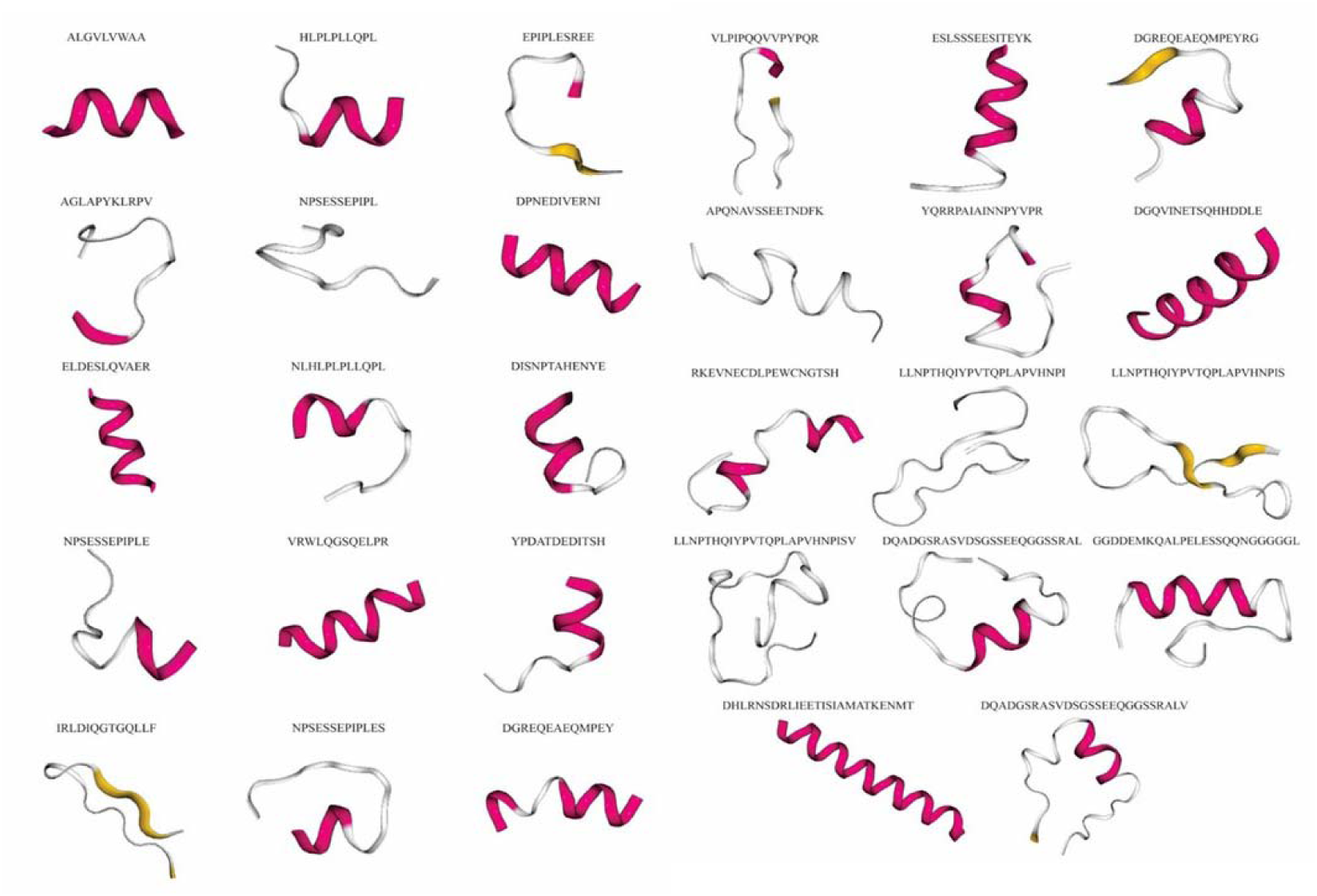
Structures of endogenous peptides in human colostrum predicted by PEPFOLD3 program. Color-coding of peptides’ secondary structure: pink, α helix; yellow, β-sheet; grey, random structure.

### 3.6. 3D structural analysis of precursor proteins

The structures of the colostrum proteins that were precursors of endogenous peptides were visualized using PyMOL software [26]. Three precursor proteins of endogenous peptides with known structures were shown: the polymeric immunoglobulin receptor protein, immunoglobulin J and lactoferrin (**Figure 4**). As evident in **Figure 4**, the peptides derived from these proteins emerged from the proteins’ cores. The peptide IRLDIQGTGQLLF, derived from the polymeric immunoglobulin receptor protein, has potential antifungal activity predicted by ClassAMP (in blue in **Figure 4**). The peptide AGLAPYKLRPV derived from lactoferrin, has two segments with secondary structure, β-sheet and α-helix, and it is potentially antiviral (in orange in **Figure 4**). The peptide DPNEDIVERNI from immunoglobulin J has one segment with β-sheet structure and is also potentially antiviral (in pink in **Figure 4**).

**Fig. 4.**
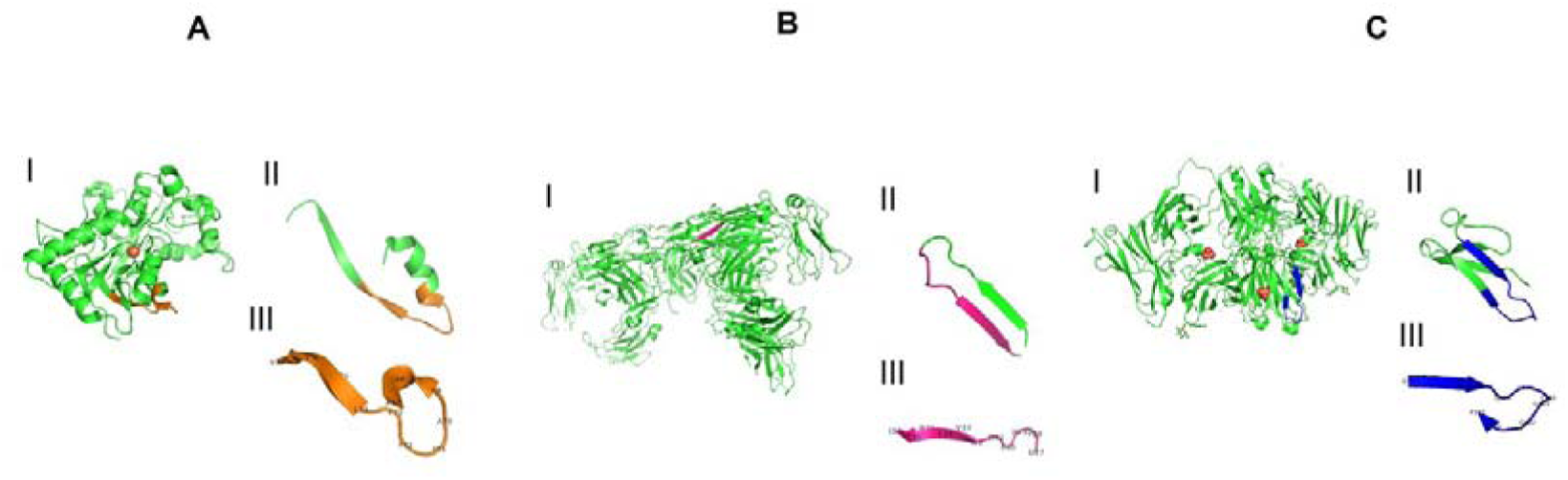
Ribbon representation of the 3D structures of three proteins identified as precursors of endogenous peptides in human colostrum. The predicted 3D structures were extracted from PDB and visualized using PyMOL software. (A) P02788, lactoferrin (PDB ID: 1H43), with the peptide AGLAPYKLRPV shown in orange; (B) P01591, immunoglobulin J (PDB ID: 6LX3), and in pink the peptide DPNEDIVERNI; (C) P01833, polymeric immunoglobulin receptor protein (PDB ID: pdb2ocw), with the peptide IRLDIQGTGQLLF shown in blue.

### 3.7. Antimicrobial activity

Endogenous peptides in human milk were able to abolish the growth of *E. coli* strain DH5alfa. Growth inhibition was achieved after exposure to peptides at 0.01 mg/mL, by 11 % with the peptide HLPLPLLQPL (from β-casein) and by 8 % with the peptide AGLAPYKLRPV (from lactoferrin) (**Supplementary Table 4**).

## 4. Discussion

Endogenous peptides result from the action of milk proteases, including plasmin, cathepsin D, and elastase. Human mammary epithelial cells secrete both activators and inhibitors of proteases that could influence the proteolytic activity in milk [6]. Previous studies on milk peptidomics have highlighted the importance of endogenous proteases in the digestion of milk proteins [6, 13,14]. In this work, sample ultrafiltration was combined with high resolution mass spectrometry and bioinformatics tools for the prospection and characterization of endogenous peptides in human colostrum with antimicrobial potential.

Twenty-nine peptide sequences were identified by high resolution mass spectrometry, which were derived from 15 precursor proteins. Several peptides identified in this work were casein-derived fragments via the activity of milk endogenous proteases. Previous studies have shown that human milk caseins are active against several microorganisms, such as *Escherichia coli*, *Helicobacter pylori*, *Listeria* spp., *Salmonella* spp., *Staphylococcus* spp., yeasts, and filamentous fungi [10]. Furthermore, we identified peptide sequences arising from the proteolysis of major proteins in milk whey, such as immunoglobulins and lactoferrin. Currently, substantial efforts are being made to find components of human milk that act in defense against emerging contagious diseases, and these efforts might prove useful in the control of such diseases, such as COVID-19 [29].

Bioinformatics analysis has been effectively used for the identification of antimicrobial peptides. In this work, four bioinformatics tools were used for the characterization of antimicrobial peptides. The Milk Bioactive Peptide Database (http://mbpdb.nws.oregonstate.edu), which includes several peptides derived from a variety of milk products, was searched. For the retrieved bioactive peptides, the AMPA tool was used to calculate the regions in the protein structure that presented potential antimicrobial activity. The potential bioactivity revealed for the endogenous colostrum peptides in this study were not previously described, to the best of our knowledge.

The APD database was searched to determine the potential antimicrobial activity [22] of the annotated peptides in human colostrum. One endogenous peptide in colostrum showed high sequence similarity (>50%) to beta-casein, which is one of the most abundant proteins in human milk. Other fragments showed 40 % similarity or lower to human proteins. However, the analysis showed that all peptides were similar to previously identified antimicrobial peptides.

When the peptide sequences were searched in the ClassAMP database, all 29 annotated sequences were predicted as antimicrobial, mostly antifungal (**Figure 2**). Antifungal peptides act specifically on fungal cells, but some may act against both fungi and bacteria [30,31]. Most studies on antimicrobial peptides emphasize antibacterial activity, rendering antifungal peptides relatively unresearched. In neonates, *Candida* spp. infection occurs in approximately 4 to 18% of ill babies and was the third most common cause of late-onset sepsis [32].

Antimicrobial peptides may present varied amino acid compositions and secondary structures, such as α-helices, β-sheets, extended helix, and loop structures [33]. For the endogenous peptides in human colostrum, the structure prediction pointed to peptides with secondary structures containing α-helix and/or random structures, seen in varied arrangements. Most frequently, the AMPs previously described have α-helix structures [33,34]; therefore, the 12 peptides containing predicted α-helix structures would be promising candidates for AMPs (**Figure 3**). However, peptides containing random structures have also been previously reported to present antimicrobial activity [34]. Therefore, the peptides NPSESSEPIPL, APQNAVSSEETNDFK, LLNPTHQIYPVTQPLAPVHNPI, and LLNPTHQIYPVTQPLAPVHNPISV could also exhibit antimicrobial properties. The three precursor proteins with known 3D structures (polymeric immunoglobulin receptor protein, immunoglobulin J and lactoferrin) revealed that the fragments coming from these proteins originated from the protein core (**Figure 4**). Therefore, we confirmed the importance of protein cleavage to expose these bioactive peptides. These peptides have secondary structures that are similar to those predicted by the Pepfold software. These structural profiles may be associated with antimicrobial activity [34].

The bioactivity of an antimicrobial peptide is strongly related to physical, chemical, and structural properties such as amphipathicity, cationicity, hydrophobicity and α- helicity [34]. On average, peptide sequences presenting a hydrophobic nature had over 65 % of hydrophobic amino acid residues in their structure (**Figure 1**). This is an important feature of AMPs as it directly influences the peptide’s ability to insert into the membranes of microorganisms [35]. Peptides with positive GRAVY values are hydrophobic, hence they favorably interact with the plasma membrane’s phospholipid bilayer, which is the first requirement for a peptide to present antimicrobial activity [36].

In all the antimicrobial peptides identified, we observed a high frequency of occurrence of glutamic acid (E), phenylalanine (F), lysine (K) and tryptophan (W) (**Fig. 1**). Additionally, the low values observed for the instability index (< 40) for five peptide sequences indicate that these peptides are fairly stable [37], and this is a further essential feature of AMPs. These sequences were derived from the polymeric immunoglobulin receptor protein, alpha S1-casein, osteopontin, vimentin and the PRA1 family protein 2. Among these sequences, we highlight the peptide IRLDIQGTGQLLF derived from the polymeric immunoglobulin receptor protein, which showed the lowest instability index (-3.83). Furthermore, the aliphatic index (AI) refers to the stability of proteins at extreme temperatures. The higher the AI, the more thermostable the protein, but peptides with an aliphatic index higher than 40 are regarded as unstable even when presenting high AI [38]. The results suggest that the endogenous antimicrobial peptides in human colostrum are thermostable, as most sequences showed a high aliphatic index (mean of 85.29). Therefore, these peptides could likely retain bioactivity at human body temperature.

Previous studies have shown that human milk can inhibit the action of several bacteria such as *S. aureus*, *E. coli*, and *P. aeruginosa*, and fungi such as those from the genus *Candida* spp. [39]. The antimicrobial activity of milk and colostrum has been attributed mainly to proteins such as immunoglobulins, lactoferrins, caseins and lysozymes. In the present study, the low inhibition of *E. coli* may be related to this strain’s resistance, as high concentrations of peptides were needed for complete inhibition. The mechanisms of action of antimicrobial peptides vary according to the target microorganism, depending on the physical properties of the membranes and the concentration and three-dimensional structure of the peptides [33]. Future *in vitro* experiments might help to propose mechanisms of action of antimicrobial peptides according to their characteristics and chemical structures.

## 6. Conclusions

Endogenous peptide sequences in human colostrum have common structural features observed in other antimicrobial peptides. From these peptides’ profile, assessed by combining nLC-Orbitrap-MS/MS and bioinformatics tools, promising sequence candidates were selected for synthesis, aiming for future confirmation of their antimicrobial activity in *in vitro* assays.

## List of abbreviations

AMP: antimicrobial peptide
COVID- 19: coronavirus disease 2019
HIV: human immunodeficiency virus
LC-HRMS: liquid-chromatography high-resolution mass-spectrometry
WHO: World Health Organization.

## CRediT Statement, author contributions: Isabele Campanhon

Validation, Formal analysis, Investigation, Writing – Original draft, Visualization; **Nicole Cavalcante**: Formal analysis, Investigation; **Rosane Nunes**: Formal analysis, Investigation; **Flávia Bezerra**: Resources, Writing – Review & editing; **Alexandre Guedes Torres**: Conceptualization, Resources, Writing – Review & editing, Supervision, Funding acquisition; **Márcia Soares**: Conceptualization, Validation, Resources, Data Curation, Writing – Review & editing, Supervision, Funding acquisition. All authors read and approved the final manuscript.

## Institutional Review Board Statement

Not applicable.

## Informed Consent Statement

Not applicable.

## Data Availability Statement

Data will be available upon request.

## Conflicts of Interest

The authors have no conflicts of interest to declare.

## Acknowledgments

The authors acknowledge the following for funding this work: Coordenação de Aperfeiçoamento de Pessoal de Nível Superior, CAPES-Brazil (Finance Code 001); Conselho Nacional de Desenvolvimento Científico e Tecnológico, CNPq-Brazil (Grant # 315579/2021-8), Fundação Carlos Chagas Filho de Amparo à Pesquisa do Estado do Rio de Janeiro, FAPERJ-Brazil (Grants # E-26/201.161/2021; E-26/010.101016/2018; E-26/010.001436/2019; and E-26/211.280/2021). I.B.C. was a recipient of a CAPES/CNPq PhD scholarship, and A.G.T. was a recipient of CNPq and FAPERJ research fellowships.

## Notes

### Competing Interest Statement

The authors have declared no competing interest.

