## Supplementary material for "Endogenous antimicrobial peptides in human colostrum: a peptidomics study by liquid-chromatography high-resolution mass-spectrometry (LC-HRMS) and bioinformatics": Supplemmental Table 1

**Supplementary Table 1.** Characteristics (mean ± SD) of the donors and their newborns during colostrum period (girls, *n*= 13 and boys, *n* = 11).

| **Characteristics (*n*= 24)** | **Mean ± SD** |
| --- | --- |
| Maternal age (years) | 26.0 ± 5.3 |
| Lactational period (days) | 2.0 ± 1.3 |
| Gestational age at birth (weeks) | 39.5 ± 1.7 |
| New-born weight at birth (kg) | 3.16 ± 0.60 |
