## Supplementary material for "Endogenous antimicrobial peptides in human colostrum: a peptidomics study by liquid-chromatography high-resolution mass-spectrometry (LC-HRMS) and bioinformatics": Supplemmental Table 2

**Supplementary Table 2.** Prediction by the ClassAMP tool of antimicrobial activity of endogenous peptides of human colostrum.

| Peptide  Sequence | ClassAMP  Database |
| --- | --- |
| ALGVLVWAA | antiviral |
| HLPLPLLQPL | antibacterial |
| EPIPLESREE | antiviral |
| AGLAPYKLRPV | antiviral |
| NPSESSEPIPL | antifungal |
| DPNEDIVERNI | antiviral |
| ELDESLQVAER | antifungal |
| NLHLPLPLLQPL | antibacterial |
| DISNPTAHENYE | antifungal |
| NPSESSEPIPLE | antiviral |
| VRWLQGSQELPR | antiviral |
| YPDATDEDITSH | antibacterial |
| IRLDIQGTGQLLF | antifungal |
| NPSESSEPIPLES | antifungal |
| DGREQEAEQMPEY | antiviral |
| VLPIPQQVVPYPQR | antifungal |
| ESLSSSEESITEYK | antifungal |
| DGREQEAEQMPEYRG | antibacterial |
| APQNAVSSEETNDFK | antiviral |
| YQRRPAIAINNPYVPR | antibacterial |
| DGQVINETSQHHDDLE | antiviral |
| RKEVNECDLPEWCNGTSH | antiviral |
| LLNPTHQIYPVTQPLAPVHNPI | antifungal |
| LLNPTHQIYPVTQPLAPVHNPIS | antifungal |
| LLNPTHQIYPVTQPLAPVHNPISV | antifungal |
| DQADGSRASVDSGSSEEQGGSSRAL | antifungal |
| GGDDEMKQALPELESSQQNGGGGGL | antiviral |
| DHLRNSDRLIEETISIAMATKENMT | antifungal |
| DQADGSRASVDSGSSEEQGGSSRALV | antifungal |
