## Supplementary material for "Endogenous antimicrobial peptides in human colostrum: a peptidomics study by liquid-chromatography high-resolution mass-spectrometry (LC-HRMS) and bioinformatics": Supplemmental Table 3

**Supplementary Table 3.** Alignment of the amino acid sequences of endogenous peptides (< 10 kDa) from human colostrum identified by mass spectrometry (nLC-MS/MS Orbitrap) recorded in the database APD.

| **Peptide sequence** | **Sequence alignment** | **Similarity (%)** | **ID APD3** | **Source/Species** | **Activity against** | **Reference** |
| --- | --- | --- | --- | --- | --- | --- |
| HLPLPLLQPL | + V **P** S **L P** + **L** V **P L** G H L **P** + **L P** L **L** Q **P L** + | 50 | AP02688 | *Streptomyces* spp. | Gram (+), Gram (-), enzime inhibitor | Kling *et al*., 2015 |
| NLHLPLPLLQPL | + **L** + **L P** I **L** G N **L L** N G L **L**  N **L** H **L P** + **L** + P **L L** + Q P **L** | 46.66 | AP00096 | European common frog, *Rana temporaria* | Gram (+) | Simmaco *et al*., 1996 |
| NPSESSEPIPL | V **P** + + + **S** L **P** L V **P L** G N **P** S E S **S** E **P** + I **P L** + | 38.46 | AP02688 | *Streptomyces* spp. | Gram (+), Gram (-), enzime inhibitor | Kling *et al*., 2015 |
| NPSESSEPIPLE | V **P** + + + **S** L **P** L V **P L** G N **P** S E S **S** E **P** + I **P L** E | 38.46 | AP02688 | *Streptomyces* spp. | Gram (+), Gram (-), enzime inhibitor | Kling *et al*., 2015 |
| EPIPLESREE | **E P** F K **L** + **S** L H L  **E P** I P **L** E **S** R E E | 40 | AP01212 | Honeybees, *Apis melífera* | Gram (+), Gram (-) | Fontana *et al*., 2004 |
| VRWLQGSQELPR | L **R** + + **Q** + **S Q** F V G S **R**  V **R** W L **Q** G **S Q** + E L P **R** | 38.46 | AP01480 | Edible fat innkeeper worm or the penis  fish, *Urechis unicinctus* | Gram (+), Gram (-), antifungal | Sung *et al*., 2008 |
| AGLAPYKLRPV | R R **L** R T T T **K L** P **P V**  A G **L** + A P Y **K L** R **P V** | 41.66 | AP02946 | ND* | Gram (-) | Oyama *et al*., 2017 |
| ELDESLQVAER | + **L** R Q **S** + **Q** F **V** G S **R**  E **L** D E **S** L **Q** + **V** A E **R** | 41.66 | AP01480 | Edible fat innkeeper worm or the penis fish, *Urechis unicinctus* | Gram (+), Gram (-), antifungal | Sung *et al*., 2008 |

*Not determined
