## Supplementary material for "Endogenous antimicrobial peptides in human colostrum: a peptidomics study by liquid-chromatography high-resolution mass-spectrometry (LC-HRMS) and bioinformatics": Supplemmental Table 4

**Supplementary Table 4.** Antimicrobial activity of endogenous peptides in human milk against *E. coli*.

| **Microorganism** | **Peptide (source protein), at 0.01mg/mL** | **Growth inhibition (%)** |
| --- | --- | --- |
| **Gram-negative**  E. coli DH5α | HLPLPLLQPL (β-casein) | 11.3 ± 2.84 |
|  | AGLAPYKLRPV (lactoferrin) | 8.40 ± 5.52 |

Growth inhibition was determined using different concentrations of the aqueous extract tested on 10^3^ cells *E. coli*, 12.5 to 150 mg/mL at 37 °C for 18 h. Next, cells were serially diluted (1/10), plated, incubated at 37 °C for 18 h and colony-forming units were counted. Values expressed as the mean ± standard deviation (*n* = 3).
